# RNF20-mediated H2B monoubiquitination protects stalled forks from degradation and promotes fork restart

**DOI:** 10.1101/2024.11.25.625131

**Authors:** Debanjali Bhattacharya, Harsh Kumar Dwivedi, Ganesh Nagaraju

## Abstract

Chromatin modifications play an important role in transcription, DNA replication and repair. Nonetheless, whether histone modifications regulate replication stress responses remains obscure. Here, we show that RNF20 localizes to and promotes H2B monoubiquitination (H2Bub) at replicating sites. Knockdown of RNF20 leads to degradation of stalled forks by MRE11 nuclease, which can be rescued by inhibition of MRE11 and co-depletion of SMARCAL1/HLTF/ZRANB3 fork remodelers. RNF20 facilitates the loading of RAD51 and RAD51C at the stalled fork sites and participates in the same pathway of RAD51/RAD51C-mediated fork protection and restart. Analyses with the RING domain and phosphorylation-deficient mutants of RNF20 showed that its catalytic activity and ATR/ATM-mediated phosphorylation are essential for its role in replication stress responses. Notably, treatment of RNF20-depleted cells with chromatin relaxing agents rescue the fork protection and restart defects. Collectively, our studies uncover the role of RNF20-mediated H2Bub in regulating the chromatin dynamics to safeguard the replicating genomes.

## Introduction

Error-free genome duplication is critical for maintaining cellular homeostasis and disease prevention (Tubbs & Nussenzweig, 2017). DNA replication is constantly threatened by obstacles arising from both endogenous and exogenous sources. Cells have evolved with multiple mechanisms to deal with replication stress to safeguard the replicating genomes. One such mechanism is that stalled forks undergo remodeling to generate reversed forks. Such reversed forks prevent the accumulation of single-stranded DNA (ssDNA) and facilitate the repair and restart of the stalled forks (Berti, Cortez et al., 2020, Cortez, 2019, Quinet, Lemacon et al., 2017). However, the reversed forks are also susceptible to degradation by various nucleases including MRE11, EXO1 and DNA2 (Pasero & Vindigni, 2017). Failure to protect stalled forks leads to the accumulation of mutations and chromosomal aberrations, eventually leading to tumorigenesis (Aguilera & Garcia-Muse, 2013, Saxena & Zou, 2022).

Homologous recombination (HR) plays an important role in the repair of DNA double-strand breaks (DSBs), thereby preventing genome instability and suppressing tumorigenesis (Kass, Moynahan et al., 2016, Nagaraju & Scully, 2007, Scully, Panday et al., 2019). RAD51 recombinase nucleates onto ssDNA at the DSB sites and promotes HR. BRCA1 and BRCA2 proteins promote RAD51 filament formation, facilitating HR (Venkitaraman, 2009, Zhao, Wiese et al., 2019). Mammalian genome encodes five RAD51 paralogs; RAD51B, RAD51C, RAD51D, XRCC2 and XRCC3 (Bhattacharya, Sahoo et al., 2022). Germline mutations in *BRCA1*, *BRCA2* and *RAD51* paralogs are known to cause breast and ovarian cancers as well as bone marrow failure syndrome Fanconi anemia (FA) (Somyajit, Subramanya et al., 2010, Somyajit, Subramanya et al., 2012, Zhao et al., 2019). RAD51 paralogs participate in the HR-mediated repair of DSBs (Nagaraju, Hartlerode et al., 2009, Nagaraju, Odate et al., 2006, Somyajit et al., 2012), activation of intra-S-phase checkpoint (Somyajit, Basavaraju et al., 2013) and mitochondrial genome maintenance (Mishra, Saxena et al., 2018). In addition to BRCA proteins, RAD51 paralogs have been implicated as recombination mediators (Somyajit et al., 2010). Studies from various groups in the past decade demonstrated that BRCA1, BRCA2 and RAD51 paralogs have repair-independent functions in protecting stalled replication forks from nucleolytic degradation (Saxena, Dixit et al., 2019, Saxena, Somyajit et al., 2018, Schlacher, Christ et al., 2011, Schlacher, Wu et al., 2012, Somyajit, Saxena et al., 2015). In addition to these proteins, RAD52, RAD54, RIF1, CtIP, FA pathway proteins such as FANCD2, FANCJ, PALB2, human CST complex, BOD1L, ABRO1, MAD2L2 and DCAF14 proteins have been shown to protect the stalled forks (Liao, Ji et al., 2018, Rickman & Smogorzewska, 2019). Nonetheless, the mechanism underlying the recruitment of these proteins to the sites of stalled forks to protect and facilitate genome duplication is unclear.

Histone posttranslational modifications (PTMs) such as ubiquitination, methylation and phosphorylation play a crucial role in gene transcription, DNA replication and repair (Chen & Tyler, 2022, Ferrand, Plessier et al., 2021, Millan-Zambrano, Burton et al., 2022). The histone H2B mono-ubiquitination (H2Bub) at K120 is primarily catalyzed by RNF20/RNF40 heterodimeric E3 ubiquitin ligase in cooperation with RAD6 E2 ubiquitin conjugase (Zhu, Zheng et al., 2005). H2B-K120ub has been shown to regulate various cellular processes, including chromatin remodeling, transcriptional activation, RNA processing and export, DNA damage response (DDR), stem cell differentiation and tissue development (Driscoll & Yan, 2023, Kato & Komatsu, 2015, Shiloh, Shema et al., 2011). Altered/loss of expression of RNF20/RNF40 has been observed in various cancer cells, including breast, lung, prostate and renal cancers, implying the role of RNF20/40 in tumor suppression (Sethi, Shanmugam et al., 2018).

Evidence from various studies showed that RNF20 participates in the repair of DSBs by HR, and this activity is dependent on K120 mono-ubiquitination of H2B by RNF20 (Moyal, Lerenthal et al., 2011, Nakamura, Kato et al., 2011, Shiloh et al., 2011, So, Ramachandran et al., 2019). Cells depleted for RNF20 or expressing H2B K120R mutant exhibited sensitivity to IR and DSB-inducing genotoxic agents MMC and CPT (Nakamura et al., 2011). These cells showed a delay in the repair kinetics monitored by γH2AX and decreased repair efficiency of DSBs by HR. Consistently, RNF20 deficient cells exhibited impaired accumulation of HR factors such as BRCA1 and RAD51 (Moyal et al., 2011, Nakamura et al., 2011, Shiloh et al., 2011). Moreover, loss of RNF20 in the mouse germ cells affects the repair of programmed DSBs and causes male infertility (Xu, Song et al., 2016). Knockdown of RNF20 in the germ cells affected chromatin relaxation and recruitment of repair factors to the sites of DSBs (Xu et al., 2016). Similarly, *Saccharomyces cerevisiae* cells lacking Bre1, an ortholog of human RNF20, showed defective repair of DSBs by HR and impaired loss of histones spanning the DSB sites (Zheng, Li et al., 2018). These studies suggest an important role of RNF20-mediated H2Bub in chromatin decompaction in facilitating HR-mediated repair. In addition to DSB repair, RNF20/Bre1 participates in DNA replication and DNA damage tolerance (DDT) pathways to facilitate error-free genome duplication (Hung, Wong et al., 2017, Liu, Yan et al., 2021, Trujillo & Osley, 2012). Bre1 localizes to chromatin, spanning the origins, and catalyzes H2Bub to promote DNA replication and nucleosome assembly (Trujillo & Osley, 2012). Localization of Bre1 to the replication-coupled damage sites promotes lesion bypass and fork recovery by HR (Hung et al., 2017, Liu et al., 2021, Northam & Trujillo, 2016). Cells lacking Bre1/RNF20 accumulate replication-associated DNA lesions and display increased micronucleation, chromosome instability and cell death (Chernikova, Razorenova et al., 2012, Hung et al., 2017, Liu et al., 2021). However, the underlying mechanism by which RNF20 participates in replication stress responses and genome maintenance is less understood.

Here, we find that RNF20 deficient cells exhibit increased replication stress markers such as micronuclei and 53BP1 nuclear bodies. RNF20 accumulates at the sites of stalled replication forks and promotes H2Bub at K120. Depletion of RNF20 leads to the degradation of stalled forks by MRE11 nuclease, which can be rescued by inhibition of MRE11 and co-depletion of fork remodelers. RNF20 knockdown affects RAD51 and RAD51 paralogs recruitment to stalled replication sites, resulting in fork degradation and impaired fork recovery. Mechanistic studies reveal that ATR/ATM-mediated phosphorylation of RNF20 and its catalytic activity are essential for replication stress responses. Strikingly, treatment of RNF20 deficient cells with chromatin relaxing agents rescue the fork protection and restart defects. Together, our studies demonstrate that RNF20-mediated H2Bub promotes chromatin decompaction at stalled fork sites and facilitates error-free genome duplication.

## Materials and methods

### Cell lines and culture conditions

Human cell lines U2OS and HeLa were grown in Dulbecco’s Modified Eagle Medium (DMEM) supplemented with 10% FBS, 1% penicillin/streptomycin (Sigma-Aldrich) and 1% Glutamax (Gibco) at 37°C in a humidified air chamber containing 5% CO2.

### Plasmids and transfections

The plasmid encoding full-length WT human RNF20 was purchased from Origene and cloned into pcDNA3β expression vector by PCR amplification. All RING domain and phosphorylation site mutants of RNF20 were generated by site-directed mutagenesis and cloned into pcDNA3β vector. The sequence of the primers used for mutagenesis has been mentioned in Supplementary Table 1. RNF20 UTR-specific shRNA was generated for this paper, and gene-specific shRNA was generated using the reported siRNA sequence and cloned into a pRS shRNA vector.

shRNF20 #1: GGGGTGAGAGCTGGAATCTCTGC

shRNF20 #2: GAAGGCAGCTGTTGAAGATTC

All other shRNA constructs used (shRAD51C, shXRCC2, shXRCC3, shRAD51, shSMARCAL1, shZRANB3 and shHLTF) were as previously reported (Dixit, Nagraj et al., 2024, Saxena et al., 2019, Saxena et al., 2018, Somyajit et al., 2013, Somyajit et al., 2015).

All plasmid transfections for transient depletion/expression were performed using a Bio-Rad gene pulsar X cell (260 V and 1050 μF). Fresh media was added to the cells 6-8 h after transfection. Cells were processed for indicated treatments/experiments 24-30 h after transfection.

### Immunoblotting

Cells were harvested and lysed in RIPA lysis buffer (50 mM Tris-HCl pH 7.5, 1% NP40, 0.5% sodium deoxycholate, 0.1% SDS, 150 mM NaCl, 2 mM EDTA, and 50 mM sodium fluoride) supplemented with cOmplete mini protease inhibitor cocktail (Roche). Protein estimation was done by standard Bradford assay. 30-50 μg of proteins were resolved on SDS-PAGE gel and transferred onto PVDF membranes (Millipore) by semi-dry transfer method (Bio-Rad Trans-Blot SD). Membranes were blocked using 5% milk in TBST (50 mM Tris-HCl, pH 8.0, 150mM NaCl, 0.1% Tween 20) and incubated with primary antibody overnight (O/N) at 4°C, followed by HRP-conjugated secondary antibody incubation for 1 h at room temperature (RT). After TBST washes, membranes were developed with chemiluminescent HRP substrate (Millipore) and imaged using Chemidoc (Bio-Rad Chemidoc Imaging System). The following primary antibodies were used in this study for western blotting: rabbit anti-RNF20 (1:2000, Abcam), mouse anti-β-ACTIN (1:2000, Santa Cruz (SC)), rabbit anti-SMARCAL1 (1:500, Abcam), rabbit anti-ZRANB3 (1:500, Abcam), mouse anti-HLTF (1:500, SC), rabbit anti-RAD51 (1:500, Abcam), mouse anti-RAD51C (1:250, SC), mouse anti-XRCC2 (1:250, SC), mouse anti-XRCC3 (1:200, SC), mouse anti-Flag (1:1000, Sigma), mouse anti-MCM3 (1:2000, SC), mouse anti-RPA70 (1:500, SC), and mouse anti-HSP70 (1:1000, SC).

### Immunofluorescence

Exponentially growing cells were seeded onto coverslips 8 h after transfection, then treated (or mock-treated) with HU as indicated. For native BrdU staining, cells were incubated with 25 μM BrdU for 48 h, washed and treated with 4 mM HU for 2 h. After treatment, the cells were washed with PBS, pre-extracted with 0.5% Triton X-100 for 90 s on ice and fixed in 4% formaldehyde for 10 min at RT. After three PBS washes, coverslips were blocked in blocking buffer (0.5% BSA and 0.5% Triton X-100 in PBS) for 30 min. The coverslips were incubated with the indicated primary antibodies for 2 h at RT. After a wash with blocking buffer, the coverslips were incubated with respective FITC/TRITC-conjugated secondary antibodies for 1 h at RT, and then stained with DAPI (1 ug/ml; Sigma-Aldrich) for 10 min before mounting onto slides with Mowiol 4-88 (Sigma). Images were acquired using an Apotome microscope (Zeiss Axio observer) and processed using ImageJ software. The following antibodies were used for performing immunofluorescence experiments in this study: rabbit anti-53BP1 (1:500, Novus Biologicals), mouse anti-Cyclin A (1:200, SC), rabbit anti-RPA70 (1:2000, Abcam), mouse anti-H2AX (pS139) (1:1000, BD Biosciences), rabbit anti-Phospho RPA32(pS4/8) (1:2000, Bethyl Laboratories), rat anti-BrdU (1:500, Abcam), mouse anti-Flag (1:1000, Sigma), rabbit anti-RAD51 (1:500, Abcam) and mouse anti-RAD51C (1:100, SC).

### Cell survival

5000 cells per well were seeded in a 24-well plate for each condition. After treatment (or mock treatment) with HU (continuous) and APH (48 h), cells were allowed to grow for 5-7 days. Later, cell survival was measured by MTT (0.3 mg/ml; Sigma-Aldrich) assay using a microplate reader (VersaMaxROM version 3.13). Percent cell survival was calculated as treated cells/untreated cells*100.

### Quantitative in-situ analysis of protein interactions at DNA replication forks (SIRF)

Exponentially growing cells were plated in 12 well plates containing coverslips the day before the experiment. Cells were incubated with 125 μM EdU for 8 min followed by HU (200 μM) or thymidine (100 μM) for 4 h and fixed with 2% PFA in PBS for 15 min at RT. Next, cells were permeabilized with 0.25% Triton X-100 in PBS for 15 min at RT. After 2x PBS washes, the Click reaction cocktail (2 mM copper sulfate, 10 μM biotin-azide, and 100 mM sodium ascorbate in PBS) was freshly prepared and added to the coverslips in a humidified chamber for 1 h at RT. After the click reaction, slides were washed with PBS and blocked with blocking buffer (10% goat serum and 0.1% Triton X-100 in PBS) for 1 h at RT followed by incubation with primary antibodies at 4°C, O/N. Mouse or rabbit anti-biotin antibody was used in conjunction with the respective antibody for the protein of interest. On the next day, PLA reactions were performed according to manufacturer’s protocol (Duolink Proximity Ligation assay, Merck). Briefly, coverslips were incubated with anti-mouse and anti-rabbit plus and minus probes in a humidified chamber for 1 h at 37°C, followed by ligation in a humidified chamber for 30 min at 37°C and finally, amplification of the annealed probes in a humidified chamber for 100 min at 37°C. Coverslips were stained with DAPI for 10 min before mounting onto slides with mounting medium (Mowiol 4-88, Sigma). Cells were imaged using an Apotome microscope (Zeiss Axio observer), and PLA foci were quantified using ImageJ software. Antibodies used for SIRF assay in this study include: rabbit anti-biotin (1:1000, CST), mouse anti-biotin (1:1500, Invitrogen), rabbit anti-RNF20 (1:500, Abcam), rabbit anti-RAD51 (1:250, Abcam), rabbit anti-RAD51C (1:100, Abcam) and mouse anti-ubiquityl histone H2B (1:250, Merck).

### DNA fiber assay

Cells were plated in a six-well plate after transfection, and after 24 h, cells were sequentially pulse-labeled with 25 µM CldU (Sigma) and 250 µM IdU (Sigma) followed by 4 mM HU treatment for 5h for fork protection assay. Alternatively, cells were pulse-labeled with 25 µM CldU and treated with 2 mM HU for 2 h to induce fork stalling. This was followed by recovery into fresh media containing 250 µM IdU for fork restart assay. Next, the cells were incubated in ice-cold PBS on ice for 10 mins, harvested, counted and re-suspended in 250 μl PBS. 3 ul of the cell mixture was mixed with 7 ul of lysis buffer on glass slides (ThermoScientific superfrost) and allowed to stand for 7 min. Slides were inclined at a 45° angle to spread the suspension and fixed in methanol:acetic acid (3:1) solution at 4°C, O/N. The following day, DNA was denatured by incubating in 2.5 M HCl for 1 h and blocked with 2% BSA in 0.1% PBST solution (1X PBS and 0.1% Tween-20). Next, the slides were incubated with primary antibodies for 2.5 h and secondary antibodies for 1 h at RT. Coverslips were mounted on the slides with a Mowiol mounting medium (Sigma) and visualized using an Apotome microscope (Zeiss Axio observer). Fiber length was measured using ImageJ software from 2-3 independent experiments and P values were calculated using Prism software. Antibodies used for performing DNA fiber studies include: rat anti-BrdU for CldU (1:500, Abcam), mouse anti-BrdU for IdU (1:250, BD Biosciences), rabbit anti-mouse IgG (Alexa Fluor 488) (1:500, Abcam) and donkey anti-rat IgG (Alexa Fluor 594) (1:500, Abcam).

### Co-immunoprecipitation

5*10^^6^ cells per condition were harvested after indicated treatments and lysed in RIPA lysis buffer (without SDS) containing cOmplete mini protease inhibitor and PhosSTOP phosphatase inhibitor cocktail (Roche). 2 mg protein from each sample was either incubated with no antibody or rabbit polyclonal anti-RNF20 antibody O/N at 4°C under rotatory agitation. The following day, protein G beads were washed multiple times in lysis buffer and incubated with cell lysate+ antibody mixture for 4 h at 4°C under rotatory agitation. Following incubation, the beads were washed 3 times in lysis buffer and protein was eluted by boiling the beads in 2x Laemmli buffer for 15 min. Western blotting (WB) was performed as described in the previous section. Antibodies used for performing immunoprecipitation and western blotting include: rabbit anti-RNF20 (1.5 μg for IP; 1:2000 for WB, ABCAM), rabbit anti-(pS/pT)Q antibody (1:500, CST), mouse anti-RPA70 (1:1000, SC) and mouse anti-β-ACTIN (1:2000, SC).

### Comet assay

Frosted glass slides (Bluestar) were coated 1% agarose at least 24 h before the experiment. Cells were treated with HU, as mentioned and harvested in PBS. After centrifugation, 30 ul of cell suspension was mixed with 270 ul of 0.5% low melting point agarose and 100 ul of this mixture was spread onto pre-coated slides. Slides were incubated in chilled lysis buffer (2.5 M NaCl, 0.1 M EDTA, 10 mM Tris-HCl pH 8, 1% Triton X-100 and 10% DMSO) O/N at 4°C. The following day, the slides were washed in electrophoresis buffer (300 mM sodium hydroxide and 1 mM EDTA, pH>13) and transferred to an electrophoresis tank filled with chilled electrophoresis buffer. The electrophoresis was performed at 1 V/cm for 20 min at RT. Slides were then washed with PBS, fixed in methanol for 5 mins at RT, washed with double-distilled H_2_O and transferred to 70% ethanol, followed by 100% ethanol for 15 min each at RT. Slides were then air-dried and stained with PI (2 μg/ml in Milli-Q). Images were acquired using an Apotome microscope (Zeiss Axio observer). Comet tail moment was measured using OpenComet plug-in in ImageJ software.

### Metaphase spreads

shRNA transfected cells were treated with 2 mM HU for 4 h along with 0.1 µg/mL colcemid (KaryoMAX, Gibco) 30 h post-transfection. Cells were then harvested and re-suspended in hypotonic solution (75 mM KCl in Milli**-**Q) and incubated in 37°C waterbath for 12 min. After centrifugation at 160 x g for 10 min, cells were fixed in 5 ml of methanol: acetic acid (3:1) fixative. Fixed cells were lysed by dropping 100 μl of the cell suspension onto chilled slides, followed by incubation on a steaming water beaker (70°C-80°C) for 90 s. Slides were then air-dried and stained with Geimsa (Sigma) stain for 30 min at RT. Excess stain was washed off, and the slides were air dried. At least 50 metaphase spreads were scored from three independent experiments using Olympus BX53 microscope.

### Quantification and statistical analysis

All experiments reported were independently replicated at least three times, unless mentioned otherwise. Data represents mean ± SD/SEM from at least three independent experiments. The statistical analysis of all experiments was performed using GraphPad Prism Version 9. The statistical significance of DNA fiber experiments, unpaired immunofluorescence experiments, SIRF experiments, comet assay and metaphase spread experiments were determined by Mann-Whitney t-test. The statistical significance of grouped immunofluorescence experiments was determined by two-way ANOVA. Significance is indicated by asterisk (*p < 0.05; **p < 0.01; ***p < 0.001; ****p < 0.0001. n.s., nonsignificant.) and p < 0.05 was considered statistically significant.

## Results

### RNF20 is required for preserving genomic integrity during replication stress

Studies from yeast and mouse model systems indicate that RNF20 contributes to DNA replication, and repair of replication-associated DNA damage and fork recovery (Chernikova et al., 2012, Lin, Wu et al., 2014, Northam & Trujillo, 2016). However, the precise function of RNF20 during replication stress in mammalian cells remains obscure. To investigate this, we generated two shRNAs; one corresponding to UTR (shRNF20 #1) and one gene-specific (shRNF20 #2) and examined replication stress markers in the control and RNF20-depleted U2OS human osteosarcoma cells. Under-replicated genomic regions are marked by 53BP1 nuclear bodies in the G1 phase cells, which are passed on to daughter cells from the previous cell cycle. Interestingly, the knockdown of RNF20 led to an increase in 53BP1 nuclear bodies in U2OS cells with aphidicolin (APH) treatment, which induces replication stress by inhibiting DNA polymerase (Figures 1A, 1B and 1C). Another marker of replication stress is the appearance of micronuclei, small extra-nuclear bodies containing chromosomal fragments that are either under-replicated or mis-segregated during cell division. We observed that RNF20-depleted cells accumulated a higher percentage of micronucleated cells than control cells, and this was further increased upon APH treatment (Figures 1D and 1E). Replication stress causes uncoupling of helicase and polymerases, exposing extensive single-stranded DNA (ssDNA) stretches which are coated by the heterotrimeric ssDNA binding protein, Replication Protein A (RPA). We found that RNF20-depleted cells accumulated higher RPA70/pRPA32 S4/8 foci than control cells in response to hydroxyurea (HU) treatment (Figures 1F, 1G, S1A and S1B). Consistently, ssDNA accumulation, scored by BrdU foci, was also elevated in RNF20-depleted cells compared to the control cells (Figures S1C and S1D). Prolonged replication stress leads to the collapsing of stalled replication forks into DNA double-strand breaks (DSBs). RNF20 knockdown led to higher γH2AX foci formation, a marker for DSBs, and an increased comet tail moment when treated with HU (Figures 1H, 1I, S1E and S1F). To further assess the role of RNF20 in preserving genomic integrity, we investigated gross chromosomal abnormalities in the form of breaks, radials and gaps generated in RNF20-depleted cells under replication stress. In agreement with our previous observations, the loss of RNF20 led to increased chromosomal aberrations (Figures S1G and S1H). RNF20-depleted cells also exhibited considerable sensitivity to HU- and APH-induced replication stress (Figures 1J and 1K). These results establish the importance of RNF20 in preventing replication stress-associated catastrophes and maintaining genomic stability during replication stress.

**Figure 1.**
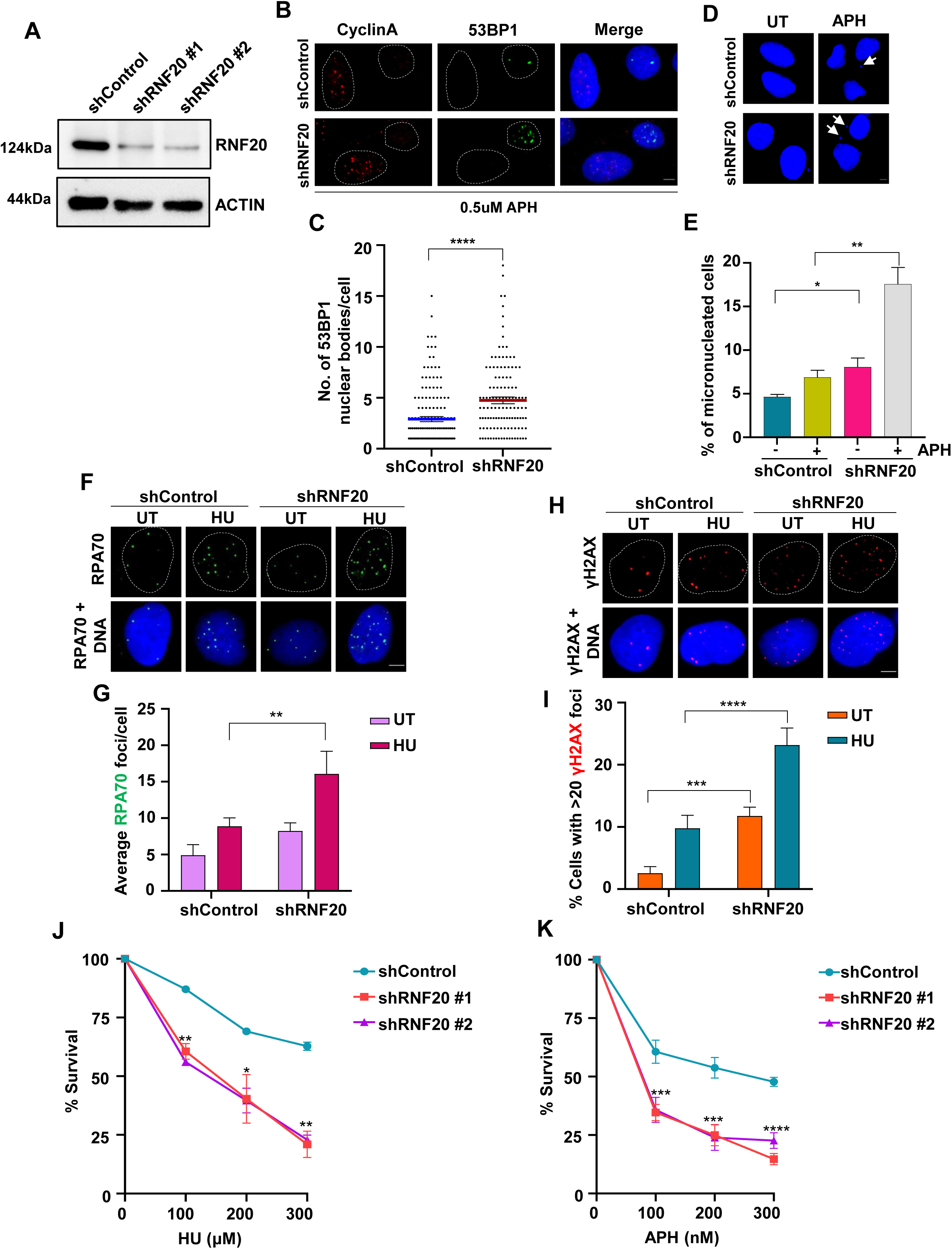
RNF20 is essential for maintaining genomic integrity during replication stress. (A) Representative immunoblot showing knockdown of RNF20 in U2OS cells using two independent shRNAs. #1 indicates a UTR specific and #2 indicates a gene-specific shRNA against RNF20. Actin serves as loading control. (B) Representative images showing 53BP1 nuclear bodies in cyclin A negative (G1 phase) U2OS cells treated with 0.5 µM APH for 12 h and the indicated shRNAs. Scale bar = 5 μM. (C) Scatter plot showing the number of 53BP1 nuclear bodies per cyclin A negative cell as shown in (B). Data represents mean ± SEM from three independent experiments. Total of ≥150 cells were analyzed for each condition. Mann-Whitney t test, *p < 0.05; **p < 0.01; ***p < 0.001; ****p < 0.0001; ns, non-significant. (D) Representative images of micronuclei formation in control and RNF20-depleted U2OS cells with or without 0.5 µM APH treatment for 12 h. Scale bar = 5 μM.(E) Bar graph showing the percentage of micronucleated cells as shown in (D). Data represents mean ± SEM from three independent experiments. Total of ≥ 300 cells were analyzed for each condition. Unpaired t test, *p < 0.05; **p < 0.01; ***p < 0.001; ****p < 0.0001; ns, non-significant. (F) Representative images of RPA70 foci formation in control and RNF20-depleted cells in untreated (UT) condition or following a recovery of 6 h after HU (4 mM, 2 h) treatment. Scale bar = 5 μM. (G) Quantification of average RPA70 foci per cell in cells as indicated in (F). Data represents mean ± SD from three independent experiments. Total of ≥ 300 cells were analyzed for each condition. Two-way ANOVA, *p < 0.05; **p < 0.01; ***p < 0.001; ****p < 0.0001; ns, non-significant. (H) Representative images showing γH2AX foci formation in control and RNF20-depleted cells in UT condition or following 6 h recovery after HU (4 mM, 2h) treatment. Scale bar = 5 μM. (I) Bar graph showing the percentage of cells with greater than 20 γH2AX foci in cells as indicated in (H). Data represents mean ± SD from three independent experiments. Total of ≥ 300 cells were analyzed for each condition. Two-way ANOVA, *p < 0.05; **p < 0.01; ***p < 0.001; ****p < 0.0001; ns, non-significant. (J and K) Plot showing survival of HeLa cells transfected with the indicated shRNAs and subjected to continuous treatment of replication stress inducing agents, HU (J) and APH (K) for 5 days. Cell survival was determined by MTT assay. Wherever not explicitly stated, shRNA #1 has been used for RNF20 knockdown.

### RNF20 localizes to and facilitates histone H2B monoubiquitination at stalled fork sites

A large-scale iPOND proteomics screen reported that RNF20 is enriched on nascent DNA at replication forks (Wessel, Mohni et al., 2019). To investigate the localization of RNF20 at replication fork sites, we performed an in-situ analysis of protein interactions at DNA replication forks (SIRF), a recently developed proximity ligation assay (PLA) system for quantitative, sensitive, and effective detection of protein association with the replication fork (Roy, Luzwick et al., 2018). Asynchronous U2OS cells were pulse labelled with a thymidine analog 5’-ethynyl-2’-deoxyuridine (EdU) for 8 mins (active forks) followed by either HU treatment (fork stalling) or thymidine chase (mature chromatin) for 4 h. Using click chemistry, a biotin moiety was conjugated to the EdU molecules. This allowed us to visualize the localization of RNF20 at the replication sites under different conditions using specific antibodies against biotin and RNF20. We indeed observed enrichment of RNF20 at active replication forks compared to the negative control and thymidine chase condition (Figures 2A and 2B). Interestingly, RNF20 SIRF signals were significantly elevated during the fork stalling condition (Figures 2A and 2B). This observation indicates that RNF20 is actively recruited to stalled replication forks.

**Figure 2.**
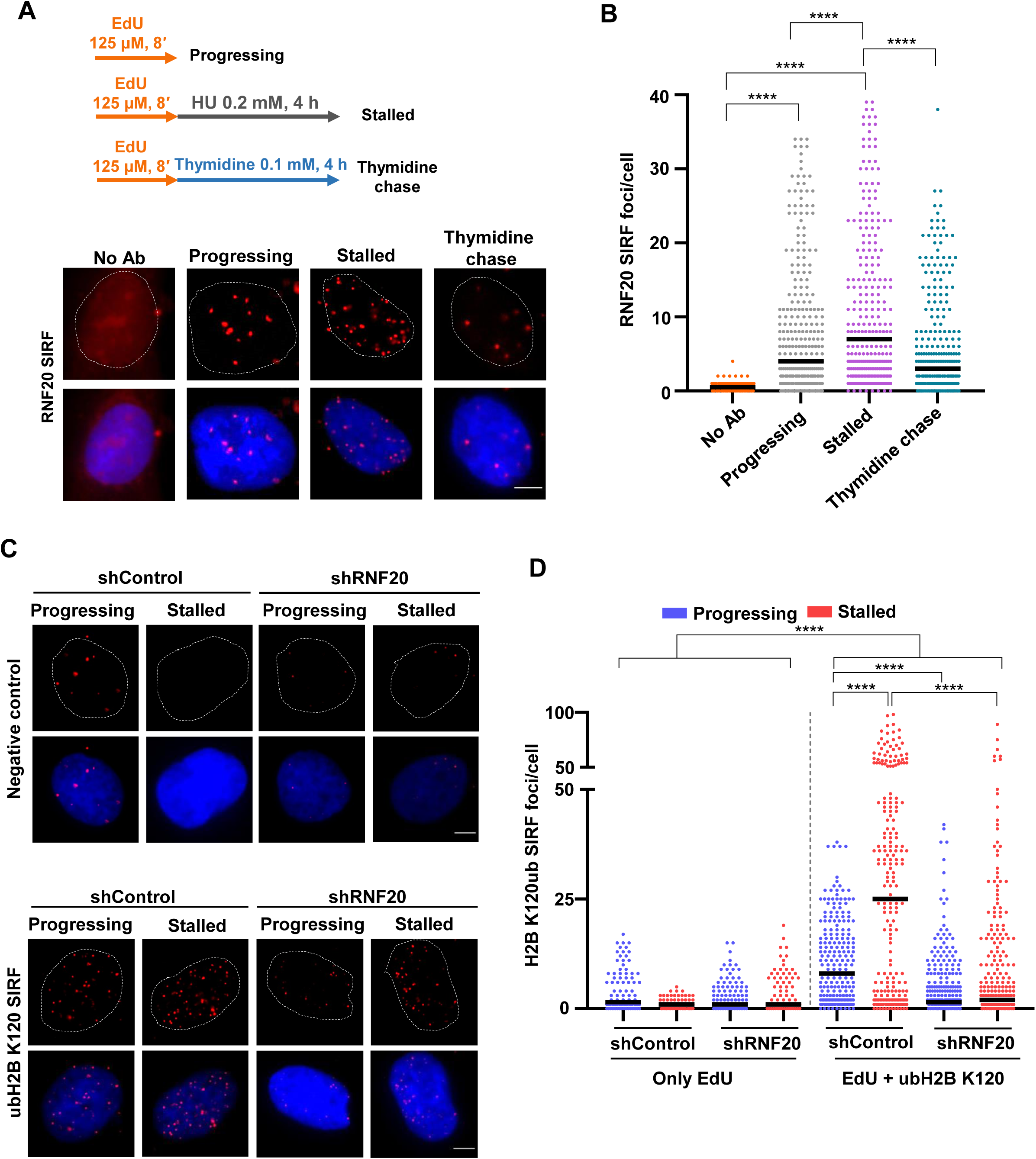
RNF20 is recruited to stalled fork sites and promotes histone H2B monoubiquitination. (A) Representative images of RNF20 SIRF signals in asynchronous U2OS cells treated with EdU ± thymidine or HU for 4 h. No Ab: no antibody is a negative control. Scale bar = 5 μM. (B) Quantification of RNF20 SIRF signals in conditions as shown in (A). Total of ≥ 300 cells were analyzed for progressing, stalled and thymidine chase conditions from three independent experiments. ≥ 150 cells were counted for no Ab. Mann-Whitney t test, *p < 0.05; **p < 0.01; ***p < 0.001; ****p < 0.0001; ns, non-significant. (C) Representative images of H2B K120ub SIRF signals at progressing and stalled replication forks in control (top) and test (bottom) set of cells transfected with shControl or shRNF20. An anti-biotin primary antibody was used for negative control whereas a mixture of anti-biotin and anti-H2B K120ub antibodies were used for test samples. Scale bar = 5 μM. (D) Quantitative scatter plot of H2B K120ub SIRF signals in conditions as shown in (C). Total of ≥ 225 cells were analyzed for each condition from three experimental repeats. Total of ≥ 100 cells were counted for the negative control samples. Mann-Whitney t test, *p < 0.05; **p < 0.01; ***p < 0.001; ****p < 0.0001; ns, non-significant.

RNF20 mono-ubiquitinates histone H2B at lysine 120 (H2B K120ub) to facilitate transcription and the recruitment of DNA damage response (DDR) proteins (Moyal et al., 2011, Nakamura et al., 2011, Pavri, Zhu et al., 2006). We investigated whether RNF20 also ubiquitinates H2B at actively progressing and stalled replication fork sites by performing SIRF in control and RNF20-depleted cells using H2B-K120ub specific antibody. H2B K120ub SIRF signals were detected at progressing replication forks, and these signals were further enriched at stalled replication forks in the control cells (Figures 2C and 2D). However, the H2B K120ub SIRF signals were abrogated at both progressing and stalled replication forks upon depletion of RNF20 (Figures 2C and 2D), indicating that H2B mono-ubiquitination at fork sites is a specific function of RNF20.

### Loss of RNF20 impairs stalled fork stability and fork recovery

The accumulation of RNF20 to stalled fork sites prompted us to investigate the function of RNF20 at stalled replication forks. We employed DNA fiber assay to analyze fork protection and restart dynamics in RNF20-depleted cells under replication stress conditions compared to control cells. U2OS cells were sequentially pulsed with 5’-chloro-2’-deoxyuridine (CldU) and 5’-iodo-2’-deoxyuridine (IdU) and subjected to prolonged replication stress by HU treatment. Reduced IdU (green) to CldU (red) tract length ratios revealed a severe fork protection defect in RNF20-depleted cells (Figures 3A and 3B). This fork protection defect was significantly rescued by treating cells with mirin (Figures 3C, 3D, and S2A), an inhibitor of MRE11 nuclease, which degrades nascent DNA at reversed forks. Fork reversal occurs through the concerted action of RAD51 and SNF2-family fork remodelers like SMARCAL1, ZRANB3 and HLTF (Bhattacharya et al., 2022, Liu, Saito et al., 2023, Taglialatela, Alvarez et al., 2017, Zellweger, Dalcher et al., 2015). Co-depletion of SMARCAL1/ZRANB3/HLTF rescued the fork protection defect in the RNF20-depleted cells (Figures 3E, 3F, S2B, S2C, and S2D). Notably, co-depletion of RNF20 with BRCA2, a major factor in the FA/BRCA pathway of fork protection, did not exacerbate the fork degradation compared to BRCA2 alone depleted cells (Figures S2E and S2F). Furthermore, RNF20-depleted cells showed a significant defect in fork restart following recovery from replication stress with a concomitant increase in stalled forks compared to the control cells (Figure 3G). However, the percentage of new origin firing events remained unaltered (Figure 3G). Together, these results clearly indicate the role of RNF20 in protecting the stalled forks from nucleolytic degradation and facilitating fork restart during replication stress.

**Figure 3.**
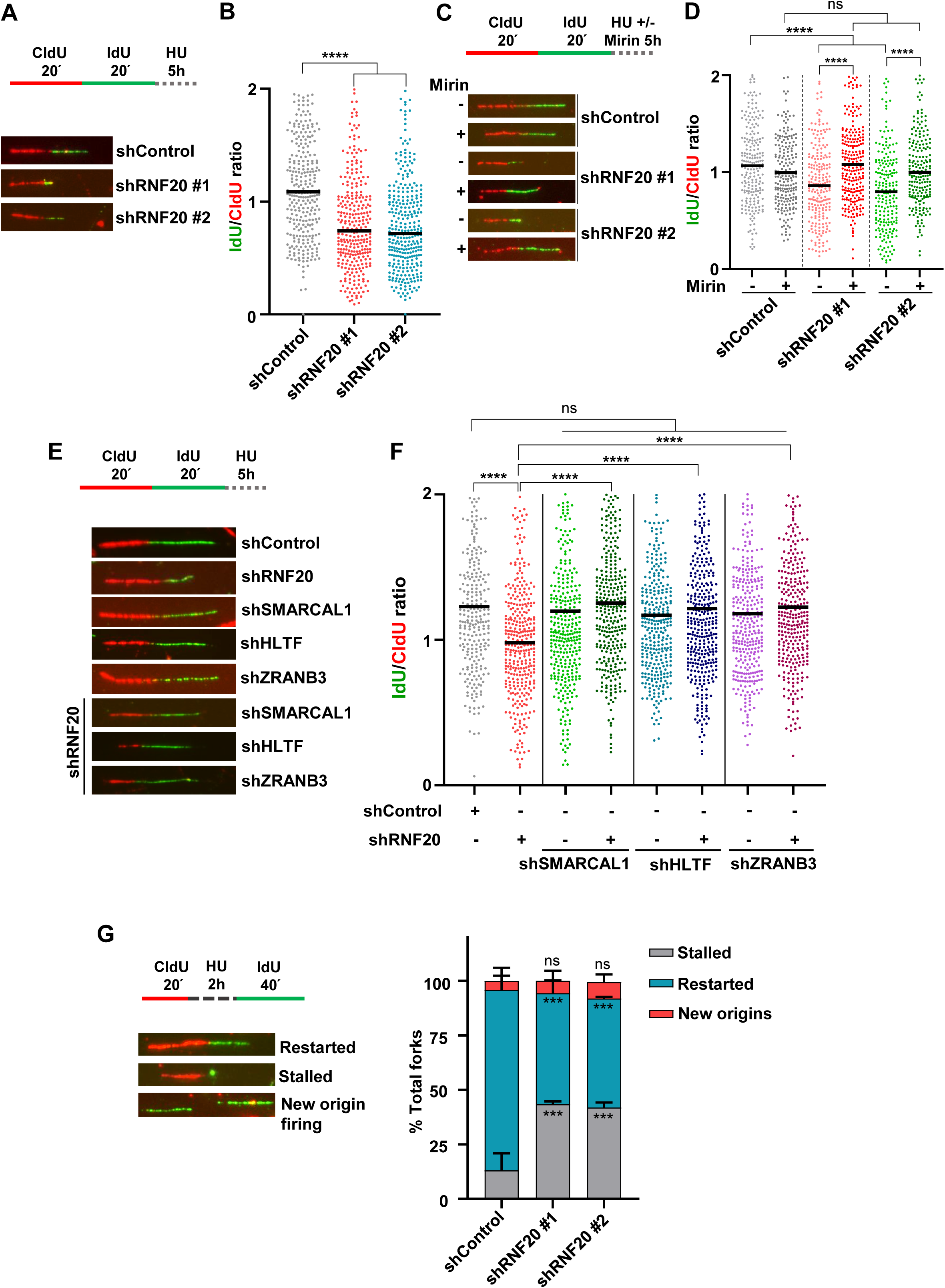
RNF20 is required for stability of stalled forks and their recovery. (A) Representative DNA fibers showing fork degradation in control and RNF20-depleted U2OS cells treated with 4 mM HU for 5h. (B) Quantification of IdU to CldU tract length ratio in cells as shown in (A). (C) Representative DNA fibers showing fork degradation in the indicated U2OS cells treated with HU ± mirin for 5h. (D) Quantification of IdU to CldU tract length ratio in cells as shown in (C). (E) Representative DNA fibers showing fork degradation in the indicated U2OS cells treated with 4 mM HU for 5h. (F) Quantification of IdU to CldU tract length ratio in cells as shown in (E). (G) Quantification of stalled, restarted forks and new origin firing events in control and RNF20-depleted cells after release from 2 mM, 2h HU treatment. Examples of various types of tracts are shown in the left panel. Stalled and restarted replication forks are shown as a percentage of all CldU-labeled tracks. DNA fiber labeling protocol has been indicated for each panel. Total of ≥ 250 fibers were analyzed for each condition from three independent experiments. Mann-Whitney t test, *p < 0.05; **p < 0.01; ***p < 0.001; ****p < 0.0001; ns, non-significant.

### RNF20 and the RAD51 paralogs participate in a common pathway of stalled fork protection and restart

Previous studies have demonstrated the roles of RAD51 paralogs (RAD51B, RAD51C, RAD51D, XRCC2 and XRCC3) in the protection and restart of stalled forks during replication stress (Guh, Lei et al., 2023, Saxena et al., 2019, Saxena et al., 2018, Somyajit et al., 2015). RAD51 paralogs exist in two major complexes – BCDX2 and CX3, and these complexes synergize with RAD51 in protecting stalled replication forks against the action of nucleolytic enzymes (Guh et al., 2023). However, the CX3 complex, but not the BCDX2 complex, exclusively participates in the fork restart (Berti, Teloni et al., 2020, Somyajit et al., 2015). Since our results showed RNF20’s role in fork protection and restart, we were curious to investigate whether RNF20 and the RAD51 paralogs perform their fork protection functions in an epistatic manner. To test this, we examined the nascent strand degradation in RNF20 and RAD51 paralogs co-depleted cells and compared it with RNF20/RAD51 paralogs alone depleted cells. Notably, while single depletions of RNF20, RAD51C, XRCC2 and XRCC3 each resulted in a significant defect in fork protection, co-depletion of RNF20 with either one of the RAD51 paralogs did not further exacerbate this effect (Figures 4A, S3A, and S3B). We next examined the status of fork restart in RNF20 and RAD51C/XRCC3 co-depleted cells by performing fork restart assay. Interestingly, the knockdown of RAD51C or XRCC3 in the background of RNF20 depletion did not alter the percentage of stalled and restarted forks when compared with RNF20/RAD51C/XRCC3 single-depletion samples (Figures 4B, S3C, and S3D). Together, these observations establish that RNF20 participates in the same pathway of RAD51 paralogs in the protection and restart of stalled replication forks during the replication stress response.

**Figure 4.**
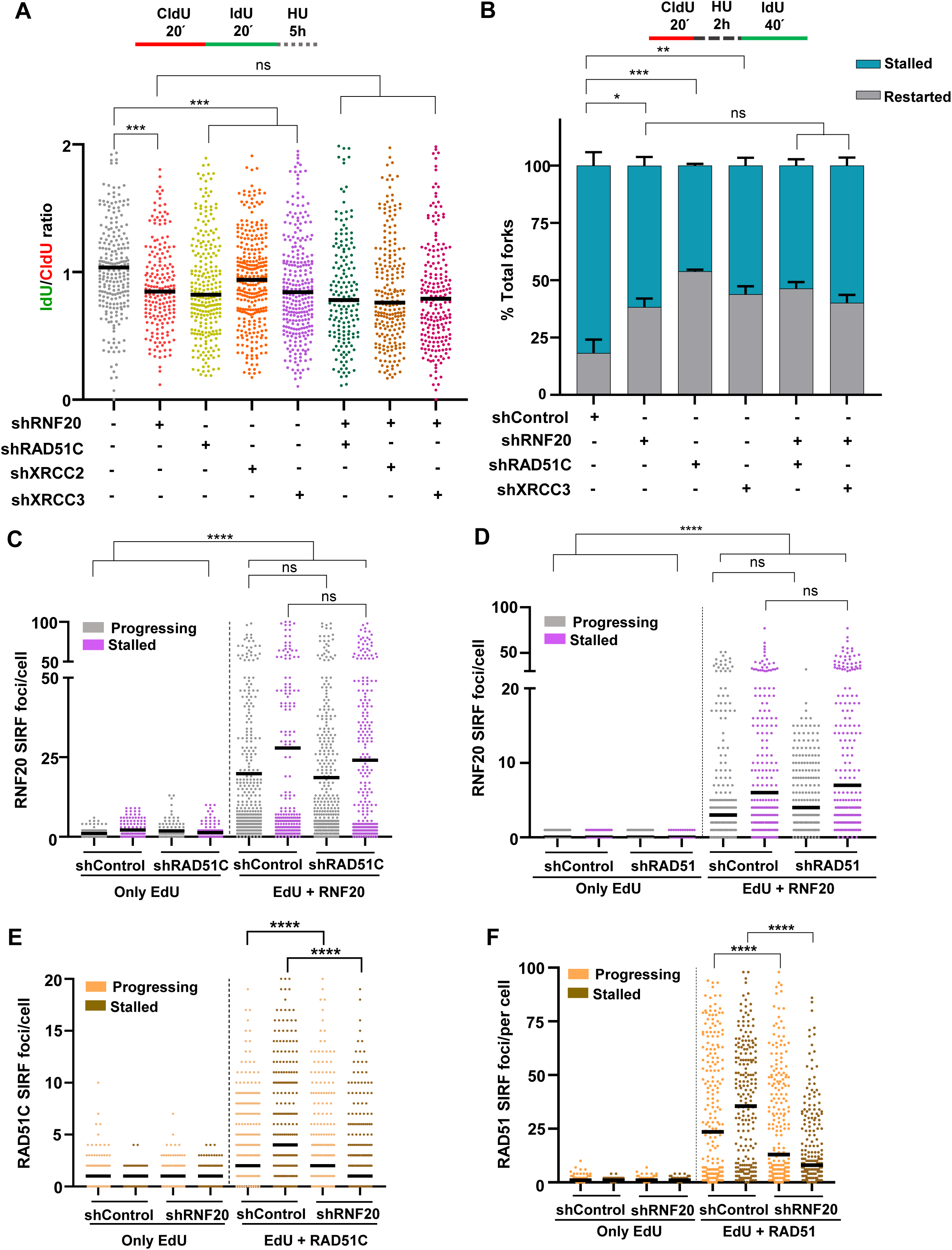
RNF20 and RAD51 paralogs function in a common pathway of stalled fork protection and restart. (A) Quantification of IdU to CldU tract length ratios in RNF20 and RAD51 paralogs depleted cells treated with 4mM HU for 5 h. Total of ≥ 200 fibers were analyzed for each condition from three independent experiments. Mann-Whitney t test, *p < 0.05; **p < 0.01; ***p < 0.001; ****p < 0.0001; ns, non-significant. (B) Quantification of the percentage of stalled and restarted forks in RNF20 and RAD51 paralogs depleted cells. Stalled and restarted replication forks are shown as a percentage of all CldU-labeled tracks. DNA fiber labeling protocol has been indicated for each panel. Total of ≥ 250 fibers were analyzed for each condition from three independent experiments. Mann-Whitney t test, *p < 0.05; **p < 0.01; ***p < 0.001; ****p < 0.0001; ns, non-significant. (C) Scatter plot of RNF20 SIRF signals at progressing and stalled replication forks in control and RAD51C-depleted cells. Total of ≥ 250 cells were analyzed for each condition. Black bars represent mean from three experimental repeats. Mann-Whitney t test, *p < 0.05; **p < 0.01; ***p < 0.001; ****p < 0.0001; ns, non-significant. (D) Quantification of RNF20 SIRF signals at progressing and stalled replication forks in control and RAD51-depleted cells. Total of ≥ 200 cells were analyzed for each condition from three independent experiments. Black bars represent median. Mann-Whitney t test, *p < 0.05; **p < 0.01; ***p < 0.001; ****p < 0.0001; ns, non-significant. (E) Scatter plot showing the number of RAD51C SIRF signals per cell in control and RNF20-deficient cells in untreated or HU-treated conditions. A total of ≥ 200 cells were analyzed for each condition from three independent experiments. Mann-Whitney t test, *p < 0.05; **p < 0.01; ***p < 0.001; ****p < 0.0001; ns, non-significant. (F) Scatter plot showing the number of RAD51 SIRF signals per cell in control and RNF20-deficient cells in untreated or HU-treated conditions. A total of ≥ 250 cells were analyzed for each condition from three independent experiments. Mann-Whitney t test, *p < 0.05; **p < 0.01; ***p < 0.001; ****p < 0.0001; ns, non-significant.

The epistatic relationship between RNF20 and the RAD51 paralogs in the replication stress responses made us ask which factor is upstream in this pathway. To decipher this, we analyzed the localization of RNF20 at progressing and stalled replication forks in RAD51C/RAD51-depleted cells by SIRF assay. Interestingly, RNF20 SIRF signals remained unchanged in RAD51C/RAD51-depleted cells compared to control cells (Figures 4C, 4D, S3E and S3F), indicating that RNF20 localization to replication fork sites is independent of RAD51C/RAD51. We verified these results with an immunofluorescence (IF) technique, where we measured the global recruitment of flag-tagged, wild-type (WT) RNF20 (expressed ectopically) to chromatin in RAD51C deficient cells. In accordance with our SIRF data, the average number of flag-RNF20 foci per cell remained unchanged between control and RAD51C-deficient cells in unperturbed or HU-treated conditions (Figures S4A and S4B).

Next, we analyzed the recruitment of RAD51C/RAD51 to replication fork sites in the absence of RNF20. Notably, the number of RAD51C/RAD51 SIRF foci at stalled replication forks was significantly reduced in RNF20-depleted cells compared to the control cells (Figures 4E, 4F, S4C and S4D). We parallelly examined the global recruitment of RAD51C/RAD51 to chromatin in RNF20-deficient cells by an IF experiment. Our results revealed a stark reduction in RAD51C/RAD51 foci per cell in HU-treated, RNF20-depleted cells compared to the control cells (S4E, S4F, S4G and S4H). Collectively, these data suggest that RNF20 functions upstream to RAD51/RAD51 paralogs during replication stress and facilitates their recruitment to stalled fork sites to promote fork protection and restart.

### E3 ubiquitin ligase activity of RNF20 is essential for its fork protection and restart function

RNF20-mediated H2BK120ub has diverse functions, including transcriptional activation and DSB repair (Moyal et al., 2011, Nakamura et al., 2011, Pavri et al., 2006). RNF20 possesses coiled-coil domains and a RING domain at the C-terminus, which is essential for E3-ubiquitin ligase activity (Figure 5A) (Foglizzo, Middleton et al., 2016). Point mutations at critical residues in the RING domain are known to abolish ionizing radiation (IR)-induced RNF20 foci formation and the interaction of RNF20 with its cognate partner, RNF40 (Moyal et al., 2011, Oliveira, Kato et al., 2014). To investigate the catalytic functions of RNF20 in replication stress responses, we mutated two critical cysteine residues in the RNF20 RING domain, C922S and C960A, by site-directed mutagenesis (Figure 5A). We depleted endogenous RNF20 in U2OS cells using a UTR-specific shRNA and simultaneously expressed the flag-tagged wild-type (WT) and RNF20 mutants. The abundance of RNF20 mutants was comparable to WT RNF20 (Figure 5B). On performing a DNA fiber based fork protection assay, we observed that the fork degradation in RNF20-deficient cells could be rescued with the expression of the WT RNF20 but not with the C922S and C960A mutants (Figures 5C and 5D). A similar result was obtained in the fork restart assay, where the percentage of stalled and restarted replication forks were rescued by expression of WT RNF20 in RNF20-depleted cells but not with the catalytically inactive mutants of RNF20 (Figures 5E and 5F). These results indicate the requirement of E3-ubiquitin ligase activity of RNF20 in the stabilization and restart of stalled replication forks. In agreement with this, H2B K120ub at stalled fork was abrogated in cells expressing RNF20 RING domain mutants compared to WT RNF20 expressing cells (Figures S5A and S5B). However, the localization of RNF20 RING domain mutants to fork sites was unaffected compared to WT RNF20 (Figures S5C and S5D). These observations further indicate that fork protection and restart defects in cells expressing catalytically inactive RNF20 are attributed to the E3-ligase activity of RNF20.

**Figure 5.**
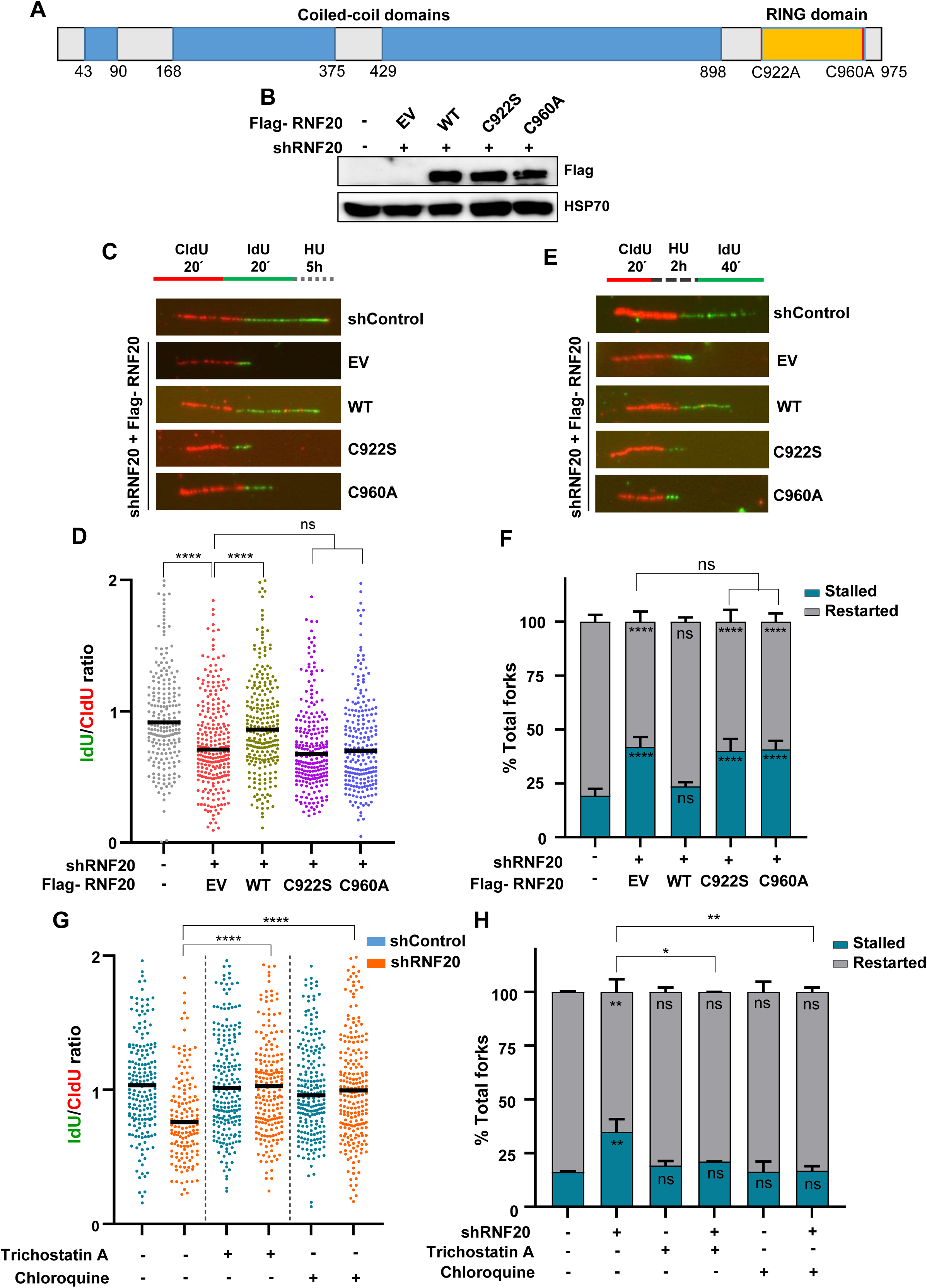
RNF20 catalytic activity is critical for fork protection and restart function. (A) Domain architecture of RNF20 E3 ubiquitin ligase. Blue boxes represent coiled-coil domains. Yellow box denotes the RING domain. Sites of point mutation in the RING domain (C922S and C960A) are indicated. (B) Representative western blot showing expression of flag-tagged wild-type (WT), C922S and C960A RNF20. HSP70 serves as the loading control. (C) DNA fiber labeling protocol (top). Representative DNA fibers showing fork protection defect in the indicated U2OS cells (bottom). (D) Quantification of IdU to CldU tract length ratio in cells as shown in (C). Total of ≥ 250 fibers were analyzed for each condition from three experimental repeats. Mann-Whitney t test, *p < 0.05; **p < 0.01; ***p < 0.001; ****p < 0.0001; ns, non-significant. (E) DNA fiber labeling protocol (top). A representative set of DNA fibres showing stalled and restarted forks in the indicated U2OS cells after release from 2 mM, 2h HU treatment (bottom). (F) Quantification of the percentage of stalled and restarted forks in cells as shown in (E). Stalled and restarted replication forks are shown as percentage of all CldU-labeled tracks. Total of ≥ 300 fibers were analyzed for each condition from four experimental repeats. Mann-Whitney t test, *p < 0.05; **p < 0.01; ***p < 0.001; ****p < 0.0001; ns, non-significant. (G) Scatter plot showing IdU/CldU ratio in the indicated cells treated with TSA (0.2 µM) or chloroquine (20 µg/ml). Total of ≥ 200 fibers were analyzed for each condition from three experimental repeats. Mann-Whitney t test, *p < 0.05; **p < 0.01; ***p < 0.001; ****p < 0.0001; ns, non-significant. (H) Bar graph showing percentage of stalled and restarted forks in the indicated cells treated with TSA (0.2 µM) or chloroquine (20 µg/ml). Stalled and restarted forks are shown as percentage of all CldU-labeled tracks. Total of ≥ 200 fibers were analyzed for each condition from three experimental repeats. Mann-Whitney t test, *p < 0.05; **p < 0.01; ***p < 0.001; ****p < 0.0001; ns, non-significant.

RNF20-mediated H2Bub has been reported to disrupt higher-order chromatin structure and cause chromatin relaxation to facilitate transcription elongation and the recruitment of different DNA repair factors during DSB repair (Fierz, Chatterjee et al., 2011, Nakamura et al., 2011, Oliveira et al., 2014). During replication stress, the regressed ends of reversed forks have been shown to undergo regular chromatinization (Schmid, Berti et al., 2018). Our data showed that RNF20-mediated H2Bub is required for RAD51 and RAD51C localization to stalled replication forks for its protection and restart. To gain insights into the mechanism of RNF20-mediated recruitment of RAD51/RAD51C to stalled fork sites, we examined whether the impaired fork protection in RNF20-knockdown cells can be rescued by nucleosome relaxation. To test this, we treated cells with chromatin-modifying agents, chloroquine and trichostatin A (TSA), which are reported to cause chromatin relaxation, and studied fork protection (Nakamura et al., 2011, Oliveira et al., 2014). Interestingly, treatment of RNF20 knockdown cells with chloroquine or TSA rescued the fork protection defect (Figures 5G, S5E, and S5F), indicating that chromatin relaxation bypasses the requirement of RNF20 in fork protection. Consistently, chloroquine or TSA also rescued fork restart defects in RNF20-deficient cells (Figures 5H, S5E and S5G). These results suggest that RNF20-mediated H2Bub is required for chromatin relaxation at stalled fork sites to recruit RAD51/RAD51C to promote fork protection and restart.

### RNF20 phosphorylation by ATR/ATM is essential for efficient fork protection and restart

ATR and ATM master kinases regulate replication stress responses and DNA damage signaling by phosphorylating numerous proteins at SQ/TQ motifs. RNF20 has been shown to undergo phosphorylation at S172 and S553 residues by ATM in response to DSBs, and this phosphorylation is required for H2Bub (Moyal et al., 2011). We wanted to investigate whether RNF20 undergoes phosphorylation in response to replication stress by ATR. To test this, we co-immunoprecipitated endogenous RNF20 from mock or HU-treated U2OS cells and performed a western blot using anti-(pS/pT)Q antibody. Interestingly, HU-induced replication stress led to a stark increase in RNF20 phosphorylation, as detected with an anti-(pS/pT)Q antibody (Figure 6A). We also observed that RPA70 interacted with RNF20 in both mock and HU-treated conditions. The lack of interaction between RNF20 and β-ACTIN served as a negative control (Figure 6A). Next, we generated point mutations at ATM-target sites in RNF20 (S172A and S553A) (Figure 6B). The cellular levels of RNF20 phosphomutants were comparable to WT RNF20 levels (Figure 6B). To examine the role of RNF20 phosphorylation, we analyzed the efficiency of RNF20 phosphomutants in protecting the stalled replication forks. Reduced IdU to CldU ratios indicated inefficient fork protection in cells expressing S172A and S553A RNF20 compared to control and WT RNF20 expressing cells (Figures 6C and 6D). We also examined the percentages of stalled and restarted forks in these cells by fork restart assay and found a similar defect in RNF20 mutant-expressing cells compared to WT cells (Figures 6E and S5H). We further analyzed the status of H2Bub at stalled replication forks in cells expressing RNF20 phosphorylation mutants by SIRF assay. Interestingly, we found reduced H2Bub-SIRF foci in RNF20-depleted cells expressing phosphorylation-deficient mutants of RNF20 compared to WT RNF20 expressing cells (Figures 6F and 6G). These results suggest that RNF20 phosphorylation is required for H2Bub at stalled fork sites, which in turn drives chromatin relaxation for replication stress responses.

**Figure 6.**
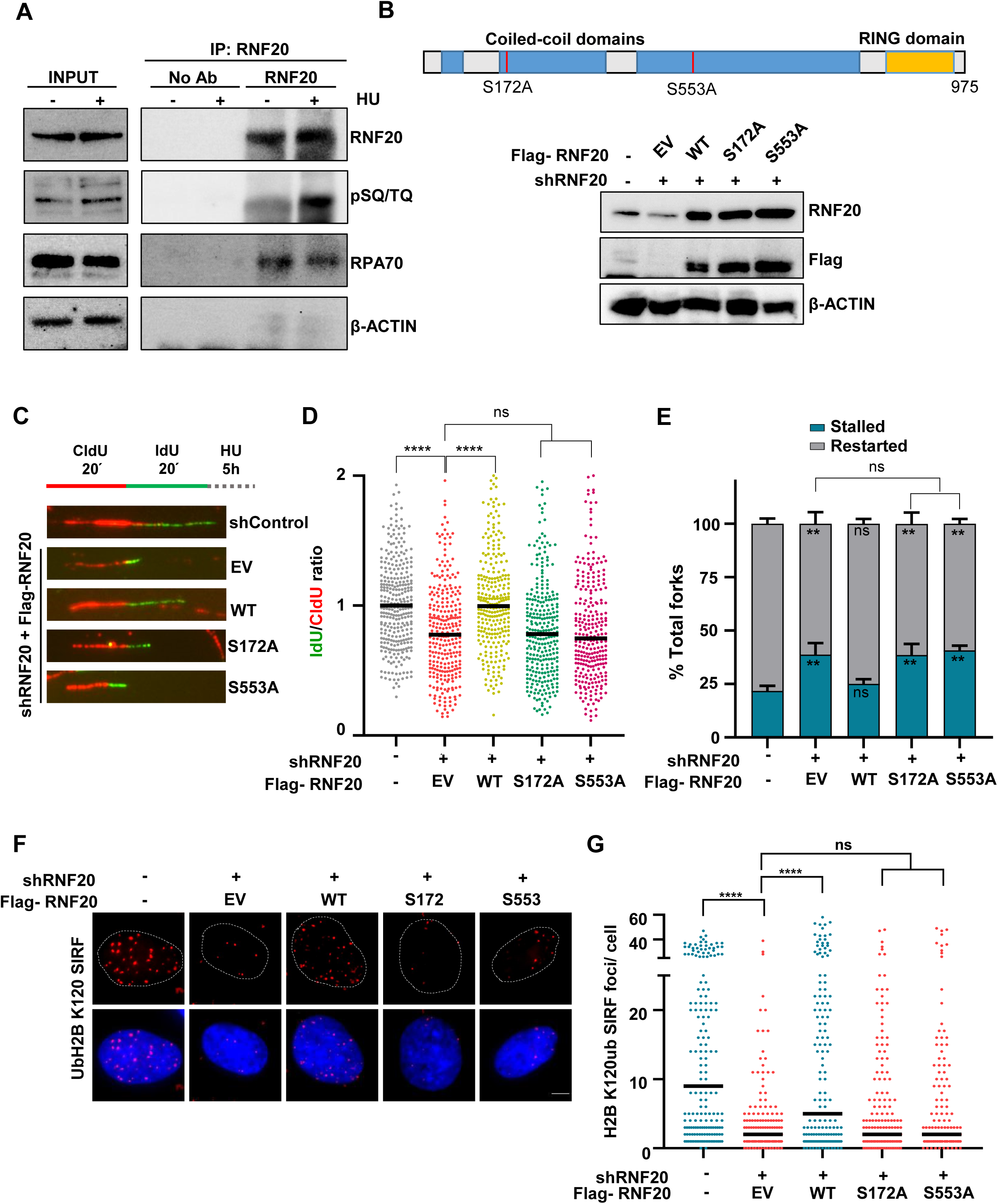
Phosphorylation of RNF20 by ATR/ATM is essential for efficient protection and restart of stalled forks. (A) Representative RNF20 co-immunoprecipitation blot (Co-IP) showing phosphorylated RNF20 (as detected by an anti-phospho-SQ/TQ antibody) in mock or HU-treated cells. RPA70 co-immunoprecipitated along with RNF20 in both UT and HU-treated conditions. β-ACTIN was taken as a negative control in the Co-IP reaction. (B) Domain architecture of RNF20 ubiquitin ligase. RNF20 phosphorylation sites and their mutants (S172A and S553A) are indicated (top). Representative immunoblot showing expression of flag-tagged wild-type (WT), S172A and S553A RNF20 constructs. Actin serves as the loading control (bottom). (C) DNA fiber labeling protocol (top). Representative DNA fibers showing the extent of fork protection in the indicated U2OS cells (bottom). (D) Quantification of IdU to CldU tract length ratio in cells as shown in (C). Total of ≥ 250 fibers were analyzed for each condition. Mann-Whitney t test, *p < 0.05; **p < 0.01; ***p < 0.001; ****p < 0.0001; ns, non-significant. (E) Quantification of percentage of stalled and restarted forks in the indicated cells. Stalled and restarted replication forks are shown as percentage of all CldU-labeled tracks. Total of ≥ 250 fibers were analyzed for each condition from three independent experiments. Mann-Whitney t test, *p < 0.05; **p < 0.01; ***p < 0.001; ****p < 0.0001; ns, non-significant. (F) Representative images depicting H2B K120ub SIRF signals in cells expressing either WT or S172A/S553A RNF20 after endogenous RNF20 depletion. (G) Quantification of H2B K120ub SIRF signals in cells as shown in (F). Total of ≥ 250 cells were analyzed for each condition. Mann-Whitney t test, *p < 0.05; **p < 0.01; ***p < 0.001; ****p < 0.0001; ns, non-significant.

## Discussion

Accurate transmission of genetic information during cell division is crucial for genome maintenance and tumor suppression. Failure to protect the stalled forks from nucleolytic degradation can lead to the accumulation of mutations, gross chromosomal rearrangements and tumorigenesis (Aguilera & Garcia-Muse, 2013, Saxena & Zou, 2022). Many HR factors, including BRCA1/2, RAD51, and RAD51 paralogs, assemble at the stalled fork sites to prevent the degradation of nascent strands and facilitate the restart of stalled forks (Bhattacharya et al., 2022, Rickman & Smogorzewska, 2019). However, the molecular mechanism by which these proteins localize to the stressed fork sites is largely unclear. RNF20-mediated H2B K120ub is important for the repair of DSBs by HR and NHEJ (Moyal et al., 2011, Nakamura et al., 2011). Data presented here demonstrates an extended role of RNF20-mediated H2Bub at K120 in regulating chromatin dynamics at the stalled fork sites to facilitate the recruitment of RAD51 and RAD51 paralogs to prevent fork degradation and promote fork restart.

Our data shows that RNF20 deficient cells are sensitive to replication stress-inducing agents, and exhibit replication stress markers such as 53BP1 nuclear bodies, micro-nucleated cells and accumulation of RPA. RNF20 has been shown to localize to DSB sites where it catalyzes H2Bub at K120 to facilitate repair of DSBs by HR and NHEJ (Moyal et al., 2011, Nakamura et al., 2011). Notably, we find that RNF20 is enriched at stalled fork sites, and RNF20-mediated H2Bub at K120 signals were also prominent at the stalled fork sites. The histone chaperone FACT has been shown to facilitate the recruitment of RNF20, BRCA1 and RAD51 to DSB sites, promoting HR-mediated DSB repair. The recruitment of RNF20 to DSB sites by FACT occurs through the SUPT16H, a component of FACT (Oliveira et al., 2014). Independent studies have also shown that RNF20/Bre1 interacts with RPA, which is essential for recruiting RNF20 to DSB sites and subsequent localization of BRCA1 and RAD51 to mediate HR repair (Li, Zhao et al., 2023, Liu et al., 2021). RPA is abundantly present at the stalled fork sites, and we find that RNF20 interacts with RPA, implying that RPA recruits RNF20 to stalled fork sites.

The localization of RNF20 to stalled fork sites is essential for protecting the forks from degradation by MRE11 nuclease and facilitating the replication restart. RAD51 and RAD51 paralogs assemble at the fork sites to prevent its degradation. RAD51 or RAD51C depletion does not affect RNF20 recruitment to fork sites. In contrast, RNF20 knockdown impairs the accumulation of RAD51 or RAD51C at the stalled replication sites, suggesting that RNF20 is upstream and facilitates the loading of RAD51 and RAD51 paralogs to fork sites to protect them from degradation. Expression of RNF20 RING domain mutants resulted in defective fork protection and its restart, implying that RNF20-mediated H2Bub is critical for fork stabilization and its recovery.

In response to DSBs, ATM kinase phosphorylates RNF20 at S172 and S553, and this phosphorylation is essential for RNF20-mediated H2Bub which in turn facilitates the recruitment of RAD51 and promotes the repair of DSBs by HR (Moyal et al., 2011). In response to replication stress, ATR kinase activates the replication checkpoint, slows down the DNA replication, and facilitates fork protection and its restart (Saldivar, Cortez et al., 2017, Simoneau & Zou, 2021). ATR regulates replication stress responses by targeting a large number of proteins. Indeed, we find that RNF20 undergoes phosphorylation, and this activation was further enhanced in response to replication stress. Strikingly, we find that RNF20-mediated fork protection and replication restart is dependent on the phosphorylation of RNF20 at S172 and S553. Importantly, H2Bub was compromised at the stalled forks in cells expressing RNF20 phosphodeficient mutants, indicating that RNF20 phosphorylation regulates the catalytic activity of RNF20.

How does RNF20-mediated H2Bub regulate replication stress response? Various studies indicate that H2Bub is required for chromatin decompaction (Kato & Komatsu, 2015). The addition of ubiquitin on H2B K120 adds a significant bulk to the nucleosome, introducing direct structural changes in chromatin, leading to its decompaction (Fierz et al., 2011). Moreover, RNF20-mediated H2Bub facilitates the recruitment of SNF2H, which belongs to the ISWI family of ATP-dependent chromatin remodelers to the sites of DSBs (Nakamura et al., 2011, Oliveira et al., 2014). Knockdown of SNF2H causes cellular hypersensitivity to DNA damaging agents and impaired accumulation of HR/NHEJ factors at DSB sites, similar to RNF20-depleted cells (Nakamura et al., 2011, Oliveira et al., 2014, Smeenk, Wiegant et al., 2013, Toiber, Erdel et al., 2013). The repair defects in RNF20/SNF2H depleted cells were rescued by chromatin relaxing agents (Nakamura et al., 2011, Oliveira et al., 2014). Using LacR/LacO heterochromatin relaxation assay, RNF20 has been shown to induce chromatin relaxation in an SNF2H-dependent manner to promote DSB repair in heterochromatin (Klement, Luijsterburg et al., 2014). These data suggest that RNF20-mediated H2Bub facilitates chromatin relaxation during transcriptional elongation and DSB repair (Klement et al., 2014, Pavri et al., 2006).

Our data shows the enrichment of RNF20 and RNF20-mediated H2Bub at the stalled fork sites, and this modification was required for the fork protection and its restart. Notably, the chromatin relaxing agents chloroquine and TSA rescued replication fork protection and restart defects in RNF20 deficient cells. These data clearly provide evidence for RNF20-mediated chromatin decompaction at stalled fork sites, which facilitates the recruitment of RAD51 and RAD51 paralogs to stalled fork sites to promote its protection and restart (Figure 7). However, further studies are required to understand whether RNF20-mediated H2Bub is essential for accumulating SNF2H or other factors at the stalled fork sites to promote chromatin decompaction. In addition to its enzymatic activity, RNF20 has been shown to serve as an adaptor to recruit BRCA1 to damage sites via TRAIP and RAP80 (Soo Lee, Jin Chung et al., 2016). Whether RNF20 similarly recruits BRCA1 or other HR factors to stalled fork sites by its physical interaction needs further investigation. H2A ubiquitination by RNF168 ubiquitin ligase is required for fork protection and restart (Schmid et al., 2018). Whether an interplay exists between RNF20 and RFN168 or RNF20 distinctly regulates replication stress responses requires further investigation.

**Figure 7.**
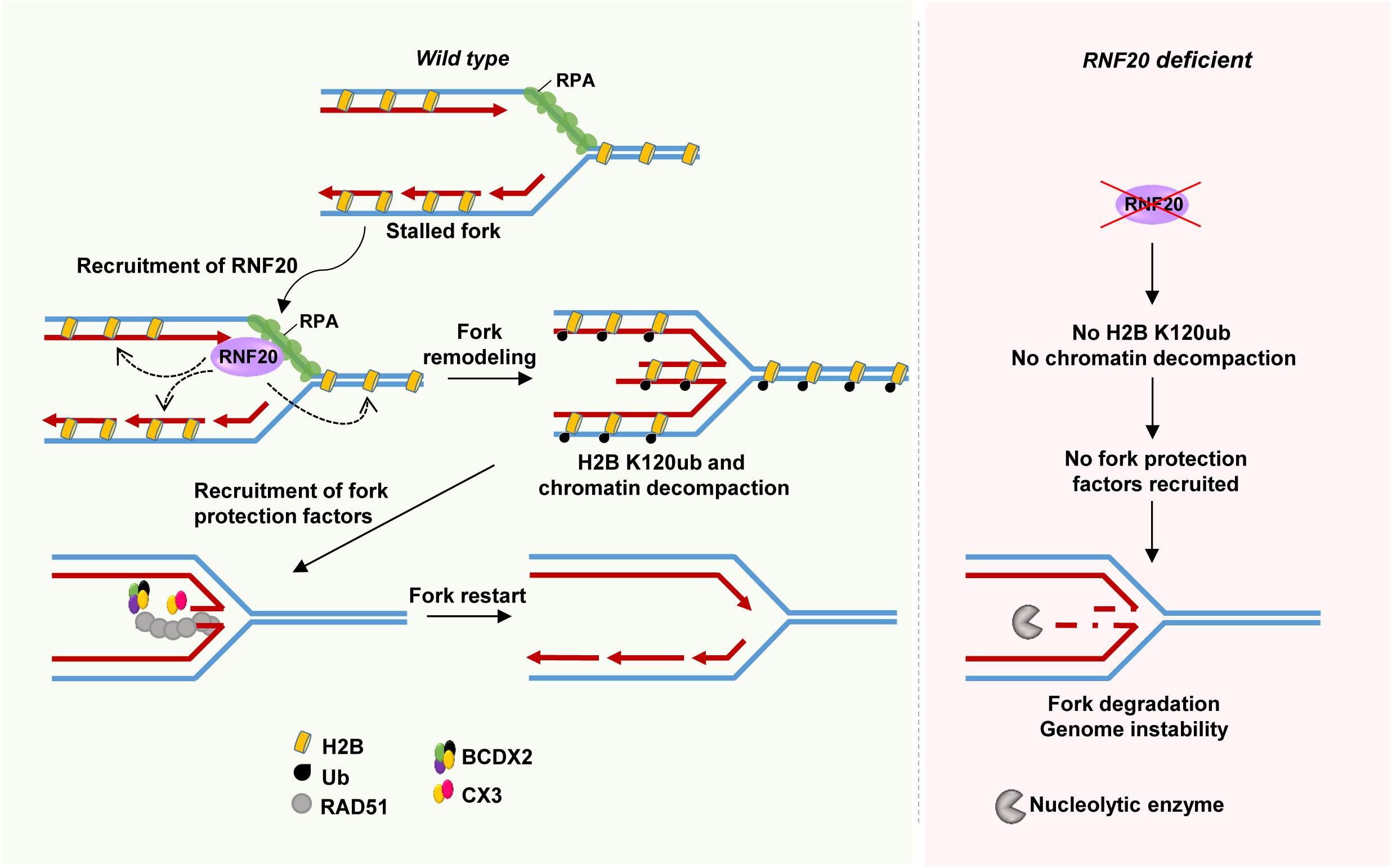
A model for RNF20-mediated H2Bub, chromatin decompaction, fork protection and restart. RPA accumulation at the stalled forks recruits RNF20 which in turn catalyze monoubiquitination of H2B K120 at the stalled replication sites. H2Bub facilitates chromatin decompaction to promote recruitment of RAD51 and RAD51 paralogs at the sites of stalled forks to safeguard the replicating genomes.

## Acknowledgements

We thank members of the GN lab for their useful discussions. We thank Dr. Kumar Somyajit for his suggestions on the manuscript. We thank Tarun Nagraj and Satyaranjan Sahoo for proofreading the manuscript. Funding by Department of Science and Technology (EMR/2015/001720; CRG/2022/003533); Department of Atomic Energy (58/14/03/2022-BRNS); Department of Biotechnology (BT/PR23498/BRB/10/1590/2017; BT/PR45508/MED/30/2414/2022); J.C. Bose fellowship (JCB/2021/000009), IISc-DBT partnership program (BT/PR27952/INF/22/212/2018) and infrastructure support provided by funding from DST and UGC are greatly acknowledged. D.B. was supported by a fellowship from the Department of Science and Technology and the Indian Institute of Science. H.K.D. was supported by a fellowship from the Indian Institute of Science.

## Author contributions

D.B. and G.N. conceived the project and designed the experiments. D.B. and H.K.D. performed the experiments. D.B., H.K.D. and G.N. analyzed the data. D.B. and G.N. wrote the manuscript.

## Conflict of interest

None

